# Reduced Expression of *Slc G*enes in the VTA and NAcc of Male Mice with Positive Fighting Experience

**DOI:** 10.1101/2020.11.10.377523

**Authors:** Dmitry A. Smagin, Vladimir N. Babenko, Irina L. Kovalenko, Anna G. Galyamina, Olga E. Redina, Natalia N. Kudryavtseva

**Affiliations:** Laboratory of Neuropathology Modeling, The FRC Institute of Cytology and Genetics SB RAS, Novosibirsk, Russia; Neurogenetics of Social Behavior Sector, The FRC Institute of Cytology and Genetics SB RAS, Novosibirsk, Russia

**Keywords:** ventral tegmental area, nucleus accumbens, prefrontal cortex, *Slc* gene family, repeated aggression, positive fighting experience

## Abstract

There are many psychiatric medications targeting the activity of SLC transporters. Therefore, further research is needed to elucidate the expression profiles of the *Slc\** genes, which may serve as markers of altered brain metabolic processes and neurotransmitter activities in psychoneurological disorders. We studied differentially expressed *Slc* genes using the transcriptomic profiles in the ventral tegmental area (VTA), nucleus accumbens (NAcc), and prefrontal cortex (PFC) of male mice with psychosis-like behavior induced by repeated aggression experience in daily agonistic interactions which are accompanied by wins. Most of differentially expressed *Slc* genes in the VTA and NAcc (12 of 17 and 25 of 26, respectively) were downregulated, which was not the case in the PFC (6 and 5, up- and down, respectively). Also, the majority of these genes were shown to have brain region-specific expression profiles. In the VTA and NAcc altered expression was observed for the genes encoding the transporters of neurotransmitters as well as inorganic and organic ions, amino acids, metals, glucose, *etc*. This means alteration in transport functions for many substrates, which results in complete disruption of all cellular and neurotransmitter processes. The neurotransmitter systems, especially, the dopaminergic one, in male mice with positive fighting experience in daily agonistic interactions undergo changes leading to profound genomic modifications which include downregulated expression of the majority of the *Slc\** genes at least in the VTA and NAcc, which is attributable to chronic stimulation of the reward systems.

## INTRODUCTION

The solute carrier (SLC) group of membrane transport proteins comprises over 400 human disease-associated genes organized into 65 families [1-6]. SLCs are responsible for transporting extremely diverse solutes, including neurotransmitters, organic molecules as well as inorganic ions, metals, *etc*. Most SLC transporters are located in the cell membrane, and some of them are located in the mitochondria or other intracellular organelles.

Several SLC transporters are targets for well-known drugs [5,7] for treatment of psychoneurological disorders. As example, inhibitors of the SLC6A* transporters (SLC6A2, SLC6A3, and SLC6A4 proteins) are promising agents for depression treatment, which act by reducing monoamines uptake in the synapses, thereby increasing their levels in the synaptic cleft. Inhibitors of the vesicular monoamine SLC18A2 carrier, which transports monoamines into synaptic vesicles, is the treatment options for Huntington disease [8], psychostimulant abuse and addiction [9]. The inhibition of SLC6A9 and SLC6A5 transporters, which regulate extracellular glycine levels in brain tissue by neuronal inhibition and excitation, is used for reducing schizophrenia symptoms, which are hypothesized to be due to a deficiency in glutamatergic signaling, because glycine also binds to excitatory NMDA receptors [10]. Therefore, altered expression of *Slc* genes (*Slc\** genes) may be indicative of altered metabolic processes and neurotransmitter systems activity in patients at risk of developing psychiatric disorders.

Our studies have shown that under repeated experience of aggression in daily agonistic interactions male mice develop a psychosis-like condition accompanied by enhanced of aggression in any social situation, irritability, disturbances of social recognition, enhanced anxiety and neurological symptoms, such as repeated stereotypies, hyperactivity, and also addiction-like state, *etc*. [11-16]. Altered metabolism and reception of excess neurotransmitters as well as changes in gene expression were found in the brains of chronically aggressive mice [17-21]. Increased expression of dopaminergic genes (*Th, Slc6a3, Snca*) against the background of activation of dopaminergic brain systems [22], was found in the ventral tegmental area [17,19], which contains dopaminergic neurons. The differentially expressed genes encoding proteins involved in the metabolism of the GABAergic and glutamatergic systems were also found in the dorsal striatum [15]. It is well known, that these brain regions are responsible for the mechanisms of positive reinforcement.

The goal of this study is to identify and analyze all *Slc\** genes that changed the expression in brain regions of male mice under repeated aggression accompanied by wins: ventral tegmental area (VTA), ventral striatum (nucleus accumbens – NAcc), and prefrontal cortex (PFC), i.e the brain regions involved in sex-, food- and drugs reward systems as well as in aggression and addiction [23-29]. We assume that differentially expressed *Slc*\* genes can serve as the markers of altered brain function in the brain regions, as possible treatment targets for psychiatric diseases, in particular, psychosis accompanied by aggression.

## METHODS AND MATERIALS

Adult male mice C57BL/6 was obtained from Animal Breeding Facility, Branch of Institute of Bioorganic Chemistry of the RAS (Pushchino, Moscow region). Animals were housed under standard conditions (12:12 hr light/dark regime starting at 8:00 am, at a constant temperature of 22+/-2°C, with food in pellets and water available *ad libitum*). Mice were weaned at three weeks of age and housed in groups of 8-10 in standard plastic cages (36×23×12cm). Experiments were performed with 10-12 week old animals. All procedures were in compliance with the European Communities Council Directive 210/63/EU on September 22, 2010. The study was approved by Scientific Council N 9 of the Institute of Cytology and Genetics SD RAS of March, 24, 2010, N 613 (Novosibirsk).

### Generation of Repeated Aggression in Male Mice

Repeated negative and positive social experience in male mice were induced by daily agonistic interactions [13,30]. Pairs of animals were each placed in a cage (28×14×10 cm) bisected by a perforated transparent partition allowing the animals to hear, see, and smell each other, but preventing physical contact. The animals were left undisturbed for two days to adapt to new housing conditions and sensory acquaintance before they were exposed to agonistic interactions. Every afternoon (14:00-17:00 p.m. local time), the cage cover was replaced by a transparent one, and 5 min later (the period necessary for animals’ activation), the partition was removed for 10 minutes to encourage agonistic interactions. The superiority of one of the mice was firmly established within two or three confrontations with the same opponent. The superior mouse (aggressive mouse, winner) would be attacking, chasing and biting another, who would be displaying only defensive behavior (withdrawal, sideways postures, upright postures, freezing or lying on the back). As a rule, aggressive interactions between males are discontinued by lowering the partition if the strong attacks has lasted 3 min (in some cases less) thereby preventing the damage of defeated mice. Each defeated mouse (loser) was exposed to the same winner for three days, while afterwards each the loser was placed, once a day after the agonistic interactions, in an unfamiliar cage with strange winner behind the partition. Each winning mouse (aggressor, winner) remained in its original cage. This procedure was performed for 20 days (once a day) and yielded an equal number of losers and winners.

Two groups of animals were analyzed in this experiment: 1) Controls – mice without a consecutive experience of agonistic interactions; 2) Winners – groups of repeatedly aggressive mice. Winners with the most expressed aggressive phenotypes (long lasting expressed aggression toward any losers every day) were selected for the transcriptome analysis. The winners 24 hours after the last agonistic interaction and the control animals were decapitated simultaneously. The brain regions were dissected by the same experimenter according to the map presented in the Allen Mouse Brain Atlas [31]. All samples were placed in RNAlater solution (Life Technologies, USA) and were stored at - 70° C until sequencing.

### Selection of Brain Regions

Transcriptomic analysis was performed in the VTA, NAcc, and PFC of male mice. The **VTA** contains 55-65% of dopaminergic cell bodies [32-34] giving rise to the dopaminergic mesolimbic and mesocortical pathways which project to the NAcc and PFC, respectively: dopamine is a major neurotransmitter that is involved in the integration of afferent signals with inhibitory or excitatory inputs [32,35,36]. It is suggested that the VTA could act as a hub converging and integrating multimodal signals toward dopaminergic systems [37]. GABAergic and glutamatergic neurons are also present in the VTA [38,39]. Most of the neurons in the **NAcc** are GABAergic medium spiny neurons (MSNs) which express D1-type or D2-type receptors [40,41]; about 1–2% are cholinergic interneurons and another 1–2% are GABAergic interneurons. GABA is the predominant neurotransmitter in the NAcc, and GABA receptors are numerous [42]. These neurons play an important role in processing reward stimuli [43]. The **PFC** is highly interconnected with other brain regions [44]. Several neurotransmitter systems are represented in the PFC, in particular dopaminergic, glutamatergic and cholinergic systems [45,46].

### RNA-Seq Analysis

The collected samples were sequenced at JSC Genoanalytica (www.genoanalytica.ru, Moscow, Russia), and the mRNA was extracted using a Dynabeads mRNA Purification Kit (Ambion, Thermo Fisher Scientific, Waltham, MA, USA). cDNA libraries were constructed using the NEBNext mRNA Library PrepReagent Set for Illumina (New England Biolabs, Ipswich, MA USA) following the manufacturer’s protocol and were subjected to Illumina sequencing. The resulting “fastq” format files were used to align all reads to the GRCm38.p3 reference genome using the TopHat aligner [47]. The Cufflinks program was used to estimate the gene expression levels in FPKM units (fragments per kilobase of transcript per million mapped reads) and subsequently identify the differentially expressed genes in the analyzed and control groups. Each brain area was considered separately for 3 vs 3 animals. Only annotated gene sequences were used in the following analysis. Genes were considered differentially expressed at *P* ≤ 0.05 and *q* < 0.05 were taken into consideration.

### Statistical Analysis

Agglomerative hierarchical clustering (AHC) was performed using XLStat Version 2016.02 software (www.xlstat.com). Pearson correlation coefficient has been used as a similarity metric for AHC analysis. The agglomeration method was unweighted pair-group average. Principal component analysis (PCA) was carried out using XLStat software. It was based on Pearson correlation metric calculated on FPKM value profiles of 48 *Slc\** genes across samples used.

## RESULTS

Differentially expressed *Slc\** genes in the VTA, NAcc and PFC of the winners are presented in Figures 1, 3, 4 and Supplement 1, Table S1.

**Figure 1.**
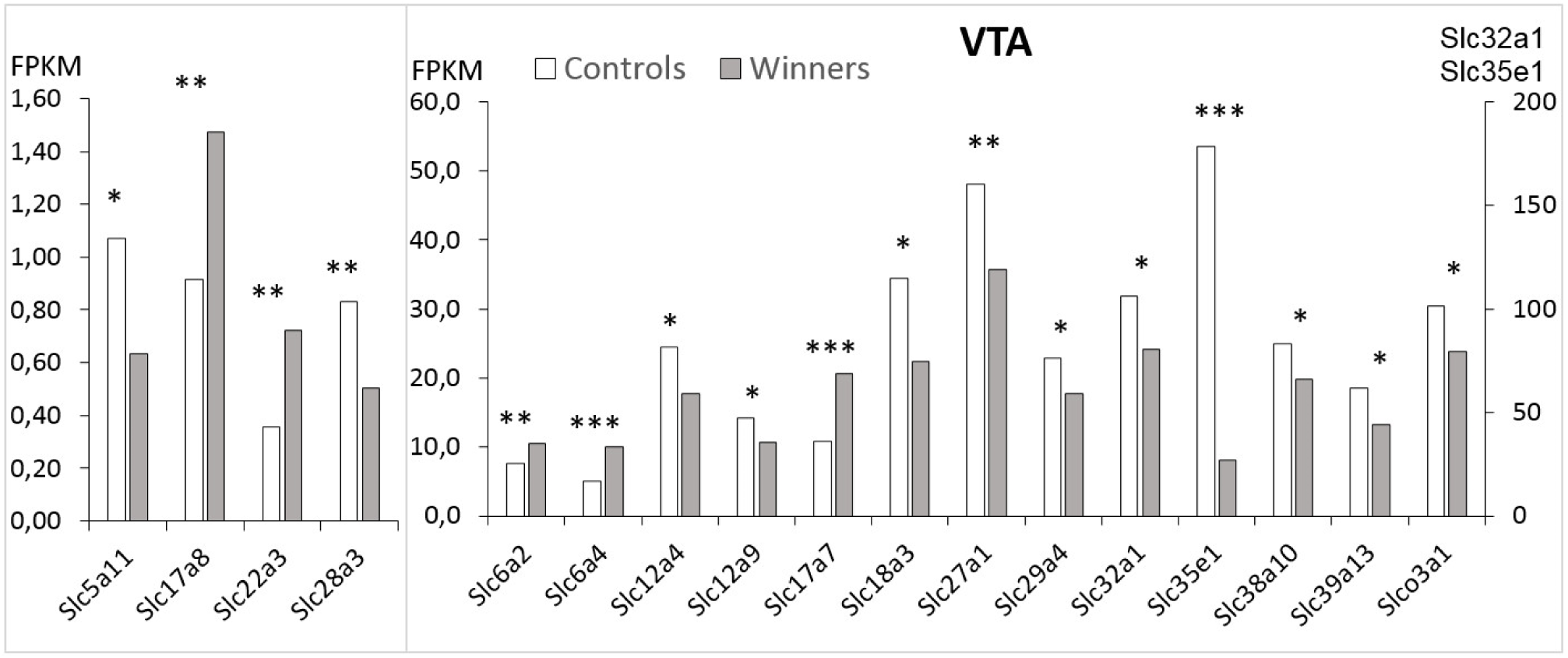
Differentially expressed *Slc\** genes in the VTA of mice. White columns – controls; grey columns – winners with repeated experience of aggression. * – *P* < 0.05; ** – *P* < 0.01; *** – *P* < 0.001. Additional statistics are shown in Supplement 1, Table 1.

### Ventral Tegmental Area

In the VTA of the winners, decreased expression compared to the controls was found for twelve *Slc* genes: *Slc5a11, Slc12a4, Slc12a9, Slc18a3, Slc27a1, Slc28a3, Slc29a4, Slc32a1, Slc35e1, Slc38a10, Slc39a13*, and *Slco3a1* genes (Figure 1). Upregulated were ***Slc6a2, Slc6a4, Slc17a7, Slc17a8***, and ***Slc22a3*** genes (Hereinafter bold indicates upregulated genes). The highest expression level as compared to the controls was found for the *Slc32a1* and *Slc35e1* genes (100-180 FPKM).

**Table 1.** Differentially expressed *Slc* genes in different brain regions according to substrate category for every gene.

|  |  |  |  |
| --- | --- | --- | --- |
| Substrate category* | The *Slc* transporters, cotransporters or symporter genes |  |  |
|  | VTA | NAcc | PFC |
| Amino acids: glycine, proline, and glutamate *etc* | ↓*Slc38a10* | ↓*Slc1a5*, ↓*Slc7a11*,<br>↓*Slc38a2*, ↓*Slc38a11* | ↑**<u>*Slc38a2*</u>** |
| Glucose, nucleotide sugars | ↓*Slc5a11*, ↓*Slc35e1* | ↓*Slc2a4* | ↓*Slc35f4* |
| Metals: zinc, magnesium, copper | ↓*Slc39a13* | ↓*Slc31a1*, ↓*Slc41a1*, | ↑***Slc39a2*** |
| Neurotransmitters: noradrenaline, serotonin, glutamate, choline, GABA, proline | ↑***Slc6a2***, ↑***Slc6a3\*\****,<br>↑***Slc6a4*** | ↓*Slc5a7*, ↓*Slc6a12*,<br>↓*Slc6a13*, ↓*Slc6a20a* | ↓*Slc6a7* |
| Vesicular transporter of neurotransmitter glutamate, acetylcholine, GABA, glycine and amino acid | ↑ ***Slc17a7***, ↑**<u>*Slc17a8*</u>**,<br>↓*Slc18a3*, ↓*Slc32a1*, | *Slc17a7* (?) | ↓<u>*Slc17a8*</u> |
| Inorganic ions: chloride, bicarbonate, hydroxide, sulfate, potassium, sodium, phosphate, monocarboxylate iodide, *etc* | ↓*Slc12a4* ↓*Slc12a9* | ↓*Slc4a2*, ↓*Slc4a5*,<br>↑***Slc4a11***, ↓*Slc5a5*<br>↓*Slc12a7*, ↓*Slc13a3*,<br>↓*Slc13a4*, ↓<u>*Slc16a12*</u> | ↓<u>*Slc5a5*</u>,<br>↑**<u>*Slc16a12*</u>**,<br>↑***Slc16a13***,<br>↓*Slc26a4* |
| Organic anions and cations, oligopeptide | ↑***Slc22a3***; ↓*Slco3a1* | ↓*Slc22a2*, ↓*Slc22a6*,<br>↓*Slc22a8*, ↓*Slco5a1* |  |
| Nucleosides | ↓*Slc28a3*, ↓ <u>*Slc29a4*</u> | ↓<u>*Slc29a4*</u> | ↑***Slc29a3*** |
| Mitochondria |  | ↓*Slc25a18* | ↑***Slc25a47*** |
| Fatty acids | ↓*Slc27a1* |  |  |
| Emphasized line - common genes for regions; **Bold** font– ↑upregulation; Regular font – ↓downregulation; * Substrate category for every gene was taken from [1-3), ** [17,19). |  |  |  |

In the VTA, differential expression of all genes indicates disturbances in transport for all substrate categories (Table 1). The differentially expressed *Slc\** genes were clustered in the VTA using AHC analysis. Figure 2 (**A)** presents the heatmap analysis based on expression profiles of the *Slc\** genes. Three distinct clusters have been identified with each of them corresponding to a specific signal transduction cascade attributable to the key genes in the brain region. AHC showed that based on highly coordinated gene expression profiles (Supplement 1, Table S2, VTA) all upregulated genes constitute one cluster: encoding the noradrenaline and serotonin transporters (***Slc6a2*** and ***Slc6a4*** genes, respectively), which carry out reuptake of monoamines into the presynaptic terminals; the ***Slc17a8*** genes encoding the protein belonging to vesicular glutamate transporter family, which mediates the uptake of glutamate into synaptic vesicles at presynaptic nerve terminals of excitatory neural cells; the ***Slc22a3*** gene encoding the protein belonging to organic cation/anion/zwitterion transporter family, which participates in dopamine transport. The second cluster was represented by a single upregulated ***Slc17a7*** gene, which demonstrates low association with other transport signaling. Additionally, the upregulation of ***Slc6a3*** gene encoding dopamine transporter was shown in the VTA [17,19].

A large cluster of 12 downregulated genes was revealed (Figure 2; Supplement 1, Table S2). In general, data indicate a deficiency in the transmembrane transport of amino acids (*Slc38a10*), inorganic (*Slc12a4, Slc12a9*), organic and oligopeptide transporters (*Slco3a1*), glucose and nucleotide-sugar transporter (*Slc5a11, Slc35e1*, respectively), fatty acids (*Slc27a1*), zinc (*Slc39a13*), the nucleoside transporters which catalyze the reuptake of monoamines into presynaptic neurons thereby regulating the intensity and duration of monoamine neural signaling, including serotonin, dopamine, *etc*. into presynaptic neurons (*Slc28a3, Slc29a4*), vesicular transporters, which are involved in acetylcholine and monoamine transport (*Slc18a3*), GABA and glycine (*Slc32a1*) *etc*. in the VTA.

**Figure 2.**
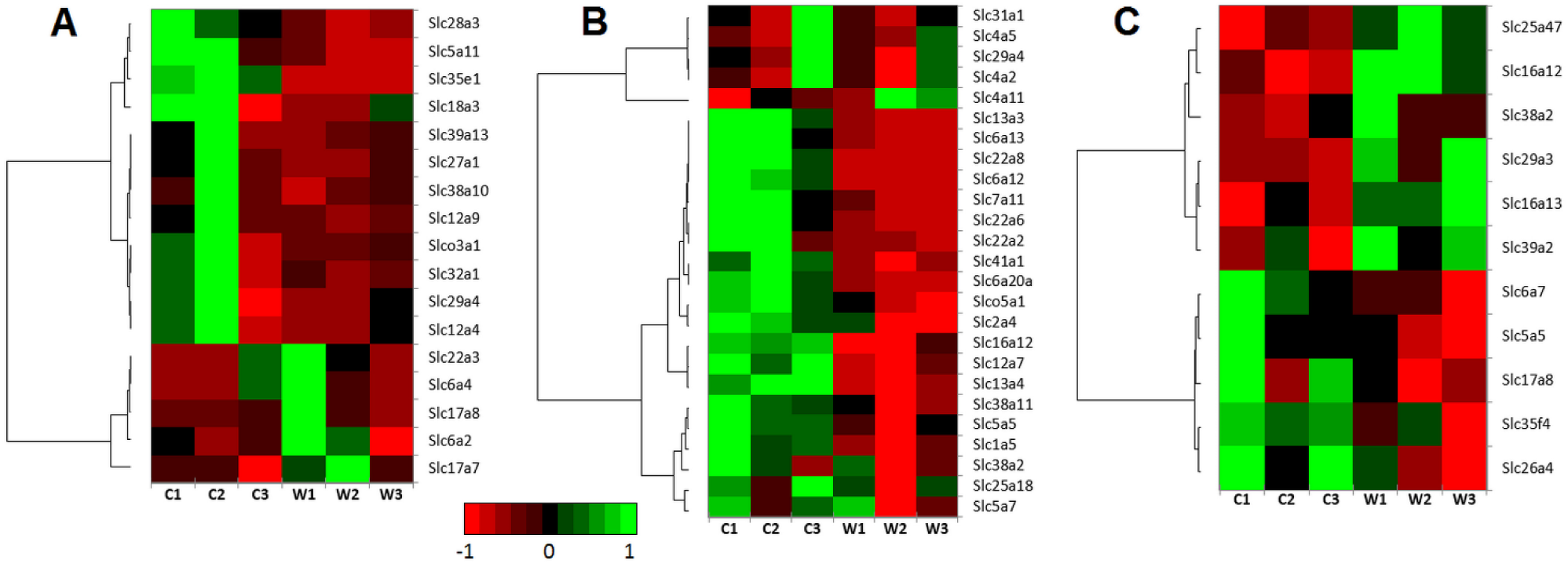
Heatmap visualization. The heatmap analysis based on expression profiles of the *Slc\** genes in the brain regions shows the expression levels of *Slc\** differentially expressed genes that were found to change in the winners (W1, W2, W3) in comparison with the control male mice (C1, C2, C3) in the VTA (A) – ventral tegmental area, NAcc (B) – nucleus accumbens, and PFC (C) – prefrontal cortex. The genes were clustered using linkage hierarchical clustering by Euclidean distance. The gene expression levels are shown with red for low, black for middle and green for high expression levels.

Each cluster corresponds to a putative specific activate signal transduction cascade. It has been confirmed that the gene clusters and samples are highly synchronized and individually correspond to the activation of neurotransmitter events according to their annotation. The largest number of positive correlations in the VTA were discovered for the expression of the *Slc12a4, Slc29a4*, and *Slco3a1* genes (Supplement 1, Table S3), which may be due to strong coordination of all processes.

### Nucleus Accumbens

In the NAcc (Figure 3, Supplement 1, Table S1) there are 26 differentially expressed genes: *Slc1a5, Slc2a4, Slc4a2, Slc4a5*, ***Slc4a11***, *Slc5a5, Slc5a7, Slc6a12, Slc6a13, Slc6a20a, Slc7a11, Slc12a7, Slc13a3, Slc13a4, Slc16a12, Slc22a2, Slc22a6, Slc22a8, Slc25a18, Slc29a4, Slc31a1, Slc38a2, Slc38a11, Slc41a1*, and *Slco5a1*. All the genes were downregulated in the NAcc, except the *Slc4a11* gene. The highest expression level was found for the *Slc25a18* gene. Expression of the *Slc17a7* gene is highly variable in the NAcc of both the controls and winners, and this gene was removed from consideration: additional statistics is needed to determine the participation of this gene in the processes in the NAcc. The heatmap analysis based on expression profiles of the *Slc\** genes in the NAcc (Figure 2, **B**) shows the expression levels of *Slc\** differentially expressed genes that were found in the winners in comparison with control male mice.

**Figure 3.**
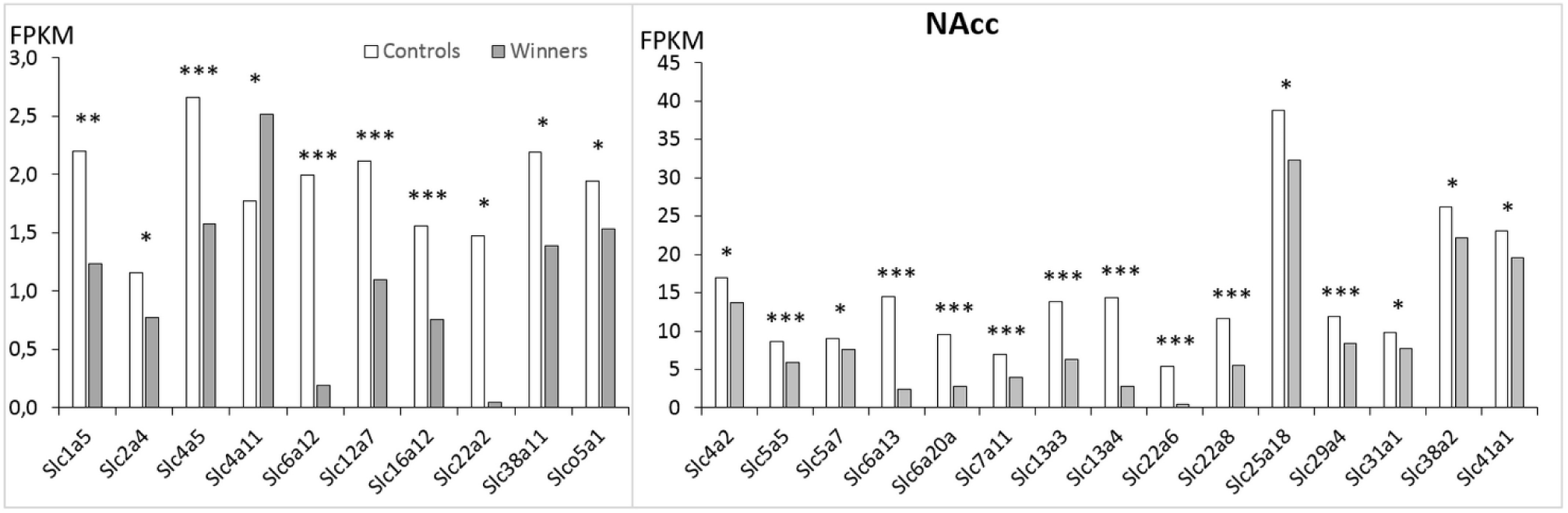
Differentially expressed *Slc\** genes in the NAcc of mice. White columns – controls, grey columns – winners with repeated experience of aggression. * – *P* < 0.05; ** – *P* < 0.01; *** – *P* < 0.001. Additional statistics are shown in Supplement 1, Table 1.

In the NAcc, downregulation of all genes (with the exception of ***Slc4a11***) indicates disturbances in transport for all substrate categories (Table 1). AHC analysis (Figure 3; Supplement 1, Table S2, NAcc) reveals 4 distinct clusters of associated genes. In the large cluster, 18 genes were highlighted. The genes in this cluster encode the carriers of neurotransmitter GABA (*Slc6a12, Slc6a13, Slc6a20a* genes), amino acids *(Slc1a5, Slc7a11, Slc38a2, Slc38a11)*, transporters of inorganic ions *(Slc12a7, Slc13a3, Slc13a4, Slc16a12 Slc41a1*, genes) and organic anions and cations (*Slc22a2, Slc22a6, Slc22a8, Slco5a1)*; as well as glucose transporters (*Slc2a4*). In the second cluster the downregulated genes encode inorganic ions (*Slc4a2, Slc4a5*), nucleoside (S*lc29a4*), and copper (*Slc31a1*) transporters. The third cluster includes the *Slc5a7* gene encoding choline, and proline carriers and mitochondrial *Slc25a18* gene. The fourth cluster includes the upregulated ***Slc4a11*** gene encoding sodium bicarbonate transporter-like protein. The *Slc29a4* gene was the only gene that changed its expression similarly in the VTA and NAcc regions.

The largest number of positive correlations with other genes were found for downregulated *Slc6a12, Slc6a13, Slc13a3* genes (Supplement 1, Table S3) encoding GABA and other neurotransmitter transporters, and for the *Slc22a8* gene encoding a protein involved in organic cation and anion exchange. The *Slc25a18* gene is expressed at the highest level.

### Prefrontal Cortex

In the PFC (Figure 4, Supplement 1, Table S1) there are 11 differentially expressed *Slc\** genes. 5 genes were downregulated: *Slc5a5, Slc6a7, Slc17a8, Slc26a4, Slc35f4*. The ***Slc16a12, Slc16a13, Slc25a47, Slc29a3, Slc38a2, Slc39a2*** genes were upregulated (Table 1). The highest expression was for the *Slc6a7* gene. The heatmap analysis based on expression profiles of the *Slc\** genes in the PFC (Figure 2, **C**) shows the expression levels of *Slc\** differentially expressed genes that were found to change in the winners in comparison with the control male mice.

**Figure 4.**
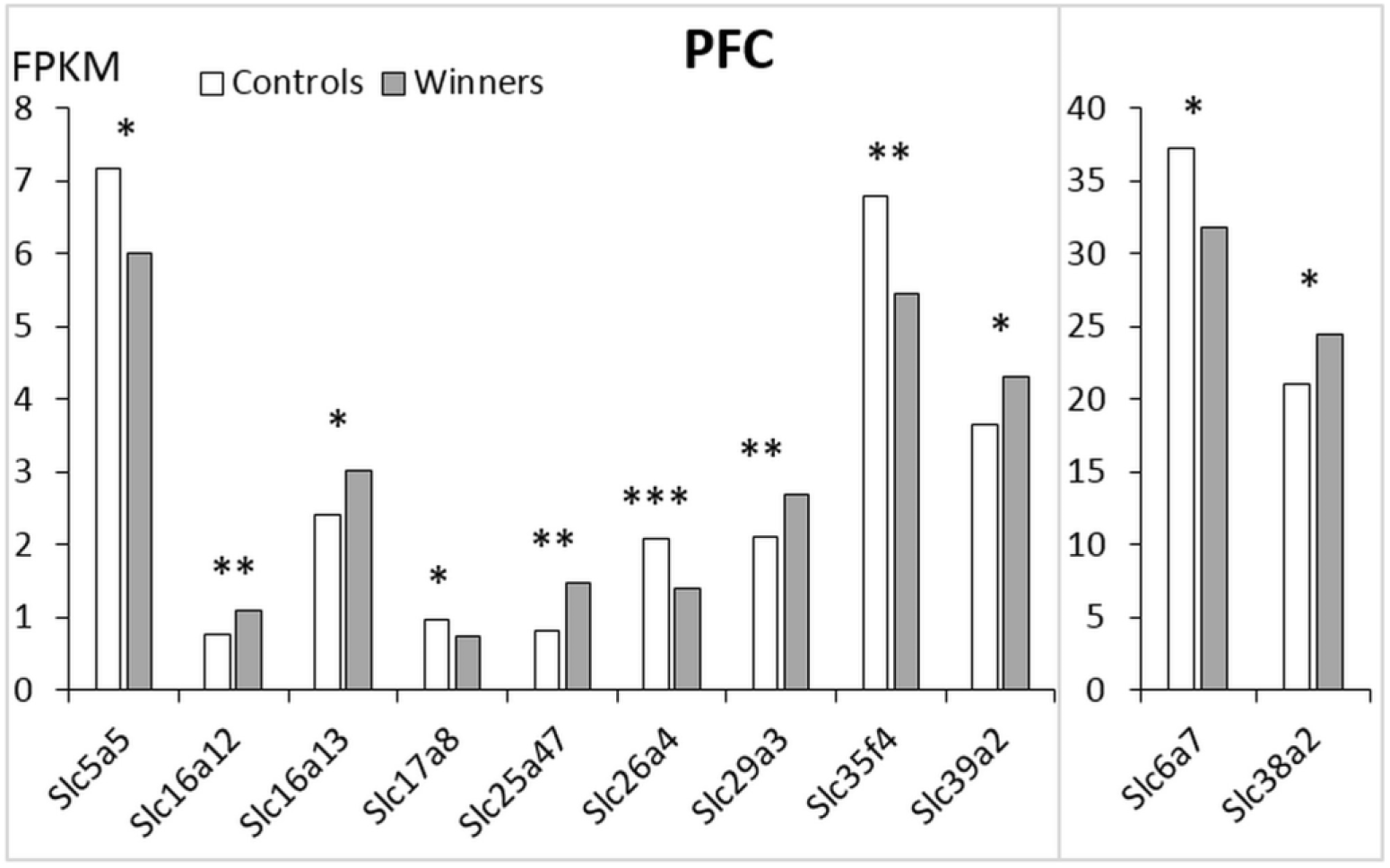
Differentially expressed *Slc\** genes in the PFC in mice. Winners – aggressive mice with repeated experience of aggression. White columns – controls, grey columns – winners. * – *p* < 0.05; ** – *p* < 0.01; *** – *p* < 0.001. Additional statistics are shown in Supplement 1, Table 1.

The AHC was performed to map expression profiles of the *Slc\** genes, which were grouped into three clusters (Supplement 1, Table S2). The first cluster of genes with reduced PFC expression contains all downregulated genes: encoding transporters of choline and proline neurotransmitters (*Slc6a7*); inorganic ions (*Slc26a4*), a putative nucleotide sugar (*Slc35f4*), and vesicular glutamate (*Slc17a8*). The second cluster is represented by a single upregulated ***Slc38a2*** gene encoding amino acid transporter. The third cluster of upregulated genes includes transporters of genes encoding monocarboxylate (***Slc16a12, Slc16a13***); mitochondrial and facilitative nucleoside ***(Slc25a47*** and ***Slc29a3***, respectively); zinc transport to the cell ***(Slc39a2***). Obviously both clusters display coordinate work of down- and upregulated genes.

The upregulated ***Slc16a13*** gene showed the largest number of correlative connections, including negative correlation with the downregulated *Slc26a4* gene as well as with downregulated *Slc5a5, Slc6a7, Slc17a8, Slc26a4, Slc35f4* (Supplement 1, Table S3). The oppositely downregulated *Slc26a4* gene correlates positively with all above-mentioned upregulated genes.

In the PFC expression of the *Slc17a7* gene is > 700 FPKM and does not differ between the experimental groups.

### PCA of *Slc\** Gene Expression

To assess the degree of brain region-specific expression of genes of interest, we performed PCA based on the covariation of 48 genes using the expression profiles of 18 samples, which comprised RNA-Seq FPKM data for three brain regions. Ovals correspond to brain regions. We observed compact clustering of samples in the VTA, NAcc, and PFC based on gene expression profiles (Figure 5 encircled). Compact clustering of samples underscores distinct pattern of considered genes expression in each brain region.

**Figure 5.**
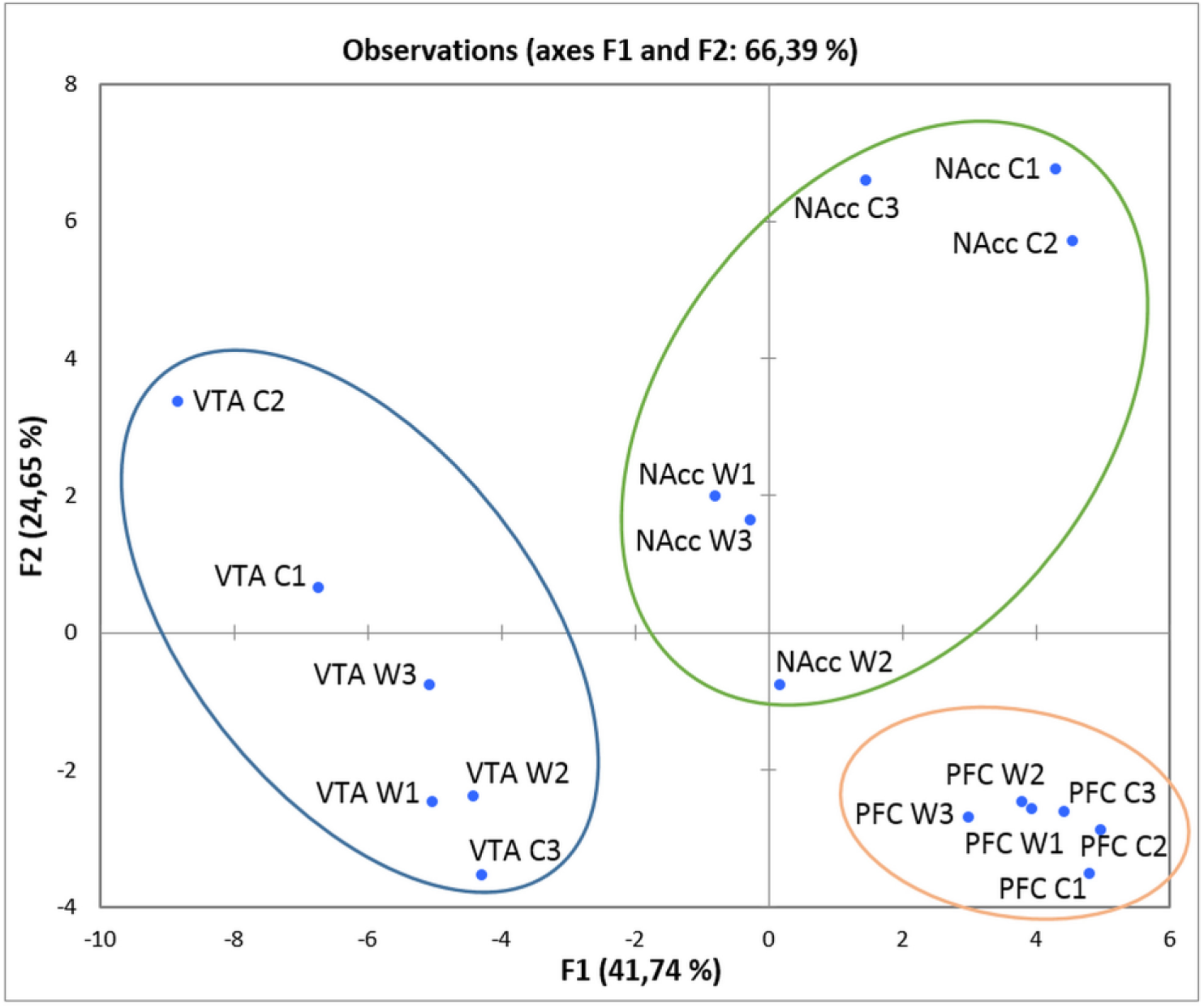
PCA plot based on covariation of genes using the expression profiles of 48 *Slc* encoding genes across 18 samples, which comprised RNA-Seq FPKM data for free brain regions. Ovals correspond to brain regions. W1, W2, W3 – winners, aggressive mice; C1, C2, C3 - control; VTA – ventral tegmental area, NAcc – nucleus accumbens; PFC – prefrontal cortex. Figure demonstrate distinct clustering of three brain regions occurred.

### Overlapping of the Differentially Expressed Genes in Brain Regions

There is overlapping of differentially expressed *Slc\** genes in different brain regions (Figure 6, Table 1). The *Slc29a4* gene was downregulated similarly in the VTA and NAcc. The *Slc17a8* gene was upregulated in the VTA and was downregulated the PFC. Similarly, changes of the *Slc38a2* and *Slc16a12* genes expression were downregulated in the NAcc and upregulated in the PFC. The *Slc5a5* similarly was downregulated in the NAcc and PFC. The other differentially expressed genes were unique for every brain region – 14 genes in the VTA, 21 genes in the NAcc and 7 genes in the PFC.

**Figure 6.**
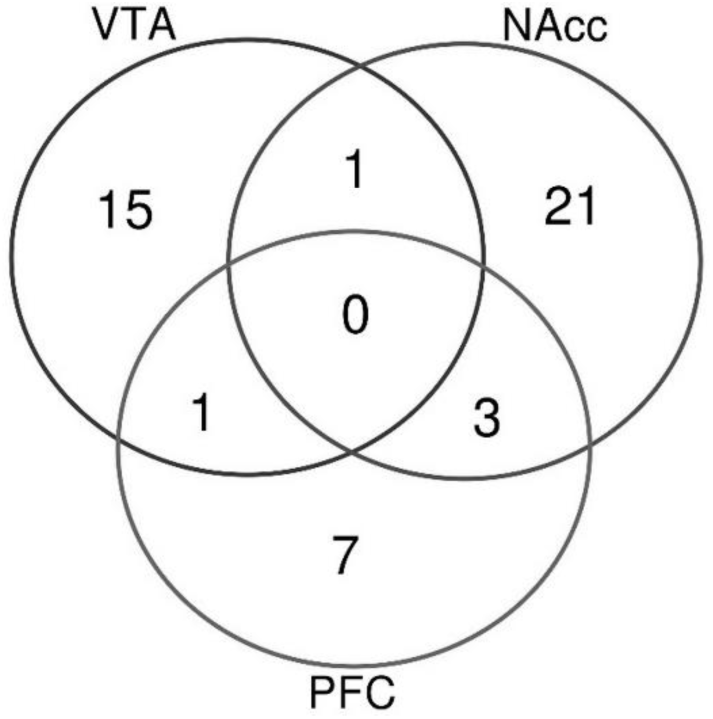
Overlapping and unique differentially expressed genes in VTA – ventral tegmental area, NAcc – nucleus accumbens; PFC – prefrontal cortex.

## DISCUSSION

It is suggested that the altered expression of the *Slc\** genes encoding the transporters can serve as a marker for altered function of substrates, including neurotransmitters [5,7,48]. Our data confirm the hypothesis that the expression level of genes encoding transporter proteins can be a measure of the activity of the corresponding transmitters [49-51]. We revealed the positive correlation between the expression of genes encoding monoamine synthesis enzymes and the expression of the *Slc6a* genes encoding the corresponding transporters in brain regions. As an example: the expression of the *Slc6a4* gene encoding the serotonin transporter significantly correlates with the expression of the *Tph2* gene encoding the rate-limiting enzyme for serotonin synthesis, and the expression of the *Slc6a2* gene encoding the noradrenaline transporter correlates with the expression of the *Dbh* gene encoding dopamine β-enzyme involved in the synthesis of noradrenaline from dopamine; the expression of the *Slc6a3* gene encoding the dopamine transporter was correlated with the expression of the *Th* gene, encoding the rate-limiting enzyme for dopamine synthesis. In addition, *Tph2* gene expression indices correlated with the *Slc6a2* and *Slc6a3* gene expression [51].

According to above data, it may be supposed that decreased expression of the *Slc\** genes encoding neurotransmitter transporters is likely to reflect a decrease of normal carrier function and, therefore, indicates an increase in the amount of substrate in the synaptic cleft and, as a consequence, activation of the corresponding neurotransmitter system. Conversely, the increased gene expression may indicate low neurotransmitter levels in the synaptic cleft leading to their decreased function. As for inorganic and organic carriers, or other carriers, changes in *Slc\** gene expression may be a consequence rather than a cause of changes in the activity of neurotransmitters in brain regions. However, based on the grounds hereinafter set forth, this simplified point of view may turn out not so self-evident.

We studied changes in expression profiles of all *Slc\** genes in the VTA, NAcc, and PFC regions associated with aggression and addiction, in male mice with psychosis-like behavioral pathology developing under long-term positive fighting experience, shown in our works [12,13,16]. Experiment revealed that the number and set of differentially expressed *Slc\** genes clearly indicate the differences in neurochemical processes in each brain region of the winners.

In the VTA, dopaminergic neurons are involved in the mechanisms of positive reinforcement of any forms of learned appetitive and motivational behaviors as well as addiction [28,52]. Disturbances in dopaminergic activity are noted in schizophrenia, Parkinson’s disease, attention deficit hyperactivity disorder [53,54], and depression [28,55,56]. It has been suggested that cellular and molecular adaptations are responsible for a sensitized dopamine activity in this brain region in response to drug abuse [57-59]. In the VTA of addicted individuals the activity of the dopamine-synthesizing enzyme tyrosine hydroxylase increases, and so does the ability of dopaminergic neurons to respond to excitatory inputs [60]. These changes may be accompanied by changes in the expression of addiction-associated dopaminergic genes. Previous experiments in animals also showed that the dopaminergic systems were activated in aggressive rats, as DA levels were elevated in the nuclei accumbens before, during and after fights [61,62]. In our experiments in mice the activation of dopaminergic systems in brain regions of the winners [22] is accompanied by increased dopaminergic *Th, Slc6a*3, and *Snca* gene expression in the VTA [17,19] accompanied by the development of addictive symptoms under positive fighting experience [16,63,64].

In this experiment upregulation of the genes encoding monoaminergic (***Slc6a2, Slc6a4*)** together with the ***Slc6a3*** gene and glutamatergic vesicular transporters (***Slc17a7, Slc17a8***) in the VTA may indicate activation of respective neurotransmitter systems in the winners in response to positive fighting experience. Interestingly, the upregulated ***Slc17a7*** gene encodes protein VGLUT1, which is used as an excitatory synapse marker. Such long-time activation may lead to a deficit of neurotransmitters in synaptic cleft and be a cause of repetitive aggressive behavior (relapse) in provoking situations. However, downregulation of genes encoding vesicular transporters of acetylcholine (*Slc18a3*), sodium-coupled nucleoside transporters (*Slc28a3, Slc29a4*), putative sodium-coupled neutral amino acid transporter (*Slc38a10*), and inhibitory vesicular amino acid GABA and glycine (*Slc32a1*) transporter indicate disturbances of the transport of substrates into synaptic vesicles, thereby exposing them to catabolism processes. We found positive correlation between the expression of genes in brain regions (Supplement 1, Table S3), which may be due to the activation of glutamatergic neurotransmission and inhibition of GABA transport to the vesicles.

Reduced expression of genes encoding glucose and nucleotide-sugar transporters (*Slc5a11* and *Slc35e1*, respectively), as well as the *Slc12a4* and *Slc12a9* genes, encoding inorganic ions transporters, and the *Slc39a13* gene encoding zinc transporter; as well as the *Slco3a1* gene encoding organic anion oligopeptide transporter were shown to indicate possible disturbances in respective functions. The largest number of positive correlations in the VTA were discovered for the expression of the *Slc12a4, Slc29a4* and *Slco3a1*, which may be due to strong coordination of all processes.

In the NAcc, most neurons are GABAergic medium spiny neurons (MSNs), which express either D1- or D2-type receptors [41], and were shown to be involved in the mechanisms of agonistic interactions in our experiments [50,65]. The number of differentially expressed genes was higher in the NAcc than in the VTA, and all genes (except *Slc4a11*) were downregulated displaying a complete disruption or even blockage of transport functions for all substrates which results in all cellular and neurotransmitter processes: glucose and sugar, neurotransmitters, amino acids, inorganic and organic transporters, *etc*.

Our results also indicate that transport processes undergo changes in the VTA, which contains the bodies of dopaminergic neurons, and in the NAcc, which is composed primarily of GABAergic MSNs [41]. GABA is one of the main neurotransmitters in the NAcc, where GABA receptors are present in large numbers. Obviously, dopaminergic inputs from the VTA, including *Slc* carriers modulate the activity of GABAergic neurons within the NAcc. The identified differentially expressed genes are as follows: *Slc29a4*, which has decreased expression both in the VTA and NAcc; *Slc17a8*, which has increased expression in the VTA and decreased expression in the PFC; *Slc38a2*, which has decreased expression in the NAcc and increased expression in the PFC. These genes are preferentially associated with the membranes of synaptic vesicles and functions of glutamate transport. Gene Ontology (GO) annotations related to these genes include L-glutamate transmembrane transporter activity and inorganic phosphate transmembrane transporter activity.

In the NAcc, the largest number of positive correlations with expression of other genes were found for the *Slc6a12* and *Slc6a13* genes encoding GABA transporters and the *Slc13a3* and *Slc22a8* genes encoding sodium-dependent decarboxylase and organic anion transporters, respectively, which may indicate that these processes are deeply intertwined.

Converging evidence from earlier [61,66-71] and more recent [29,72,73] studies have supposed that aggression is rewarding. Positive fighting experience in daily agonistic interactions leads to the activation of dopaminergic systems, in particular in the NAcc, dorsal striatum, and amygdala, as shown earlier [22,61,62] and to an increase in dopaminergic *Th, Slc6a3* and *Snca* gene expression in the VTA [17,19]. Our research demonstrates upregulation of the ***Slc6a2, Slc6a3, Slc6a4, Slc17a7, Slc17a8*** neurotransmitter genes, which may be a consequence of co-activation of monoaminergic and glutamatergic systems in the VTA, a core region in aggression and addiction circuitries.

The PFC is highly interconnected with many brain regions, including extensive connections with subcortical and other cortical structures, notably the thalamus, the basal ganglia, the hypothalamus, the amygdala, the hippocampus *etc*. Several neurotransmitter systems are represented in the prefrontal cortex, such as dopamine and cholinergic systems. It can be suggested that upregulation of the ***Slc16a12, Slc16a13, Slc25a47, Slc29a3, Slc38a2, Slc39a2*** genes and decreased expression of other genes may reflect the processes in different brain regions.

Our findings show that in male mice with psychosis-like state exhibiting aggression, altered activity of neurotransmitter systems, especially the dopaminergic one, leads to a restructuring of all metabolic and neurotransmitter processes in a way specific for each brain region. It can be assumed that in chronically aggressive male mice an overall decrease in the expression of the *Slc\** genes, mainly in the NAcc, which under positive fighting experience may be ascribed to chronic stimulation of the reward systems, over time may cause aggression relapse, as shown in our experiments in male mice under repeated experience of aggression [16,63].

When analyzing these findings, it was natural to ask: what processes lead to the downregulation of most *Slc\** genes encoding different types of substrate-specific transporter proteins in the NAcc and VTA? It is very important to glean in-depth knowledge about this phenomenon because transporters are considered to be promising therapeutic targets in the treatment of many diseases [7]. It has been shown that natural rewards, such as food (glucose) and sex, as well as pharmacological manipulations are accompanied by activation of dopaminergic systems and reduce the electrophysiological excitability of GABA-containing MSNs in the NAcc [74,75]. In our model aggression development is against the background of positive reinforcement and reduced expression of the *Slc* genes may indicate the inactivation NAcc, which is consistent with literature data that rewarding stimuli reduce the activity of NAcc MSNs, whereas aversive treatments increase the activity of these neurons because of NAcc D2-receptor inhibitory function plays a critical role in reward mechanism [74]. If so, then the key genes may be rather the genes that are responsible for electrical impulse conduction, i.e., genes encoding inorganic substrates transporters. These data should be taken into consideration to explain the overall decrease in the expression of most *Slc\** genes in the VTA and NAcc.

Thus, addiction-like state developing under positive fighting experience [12,16,63], similarly to chronic drug use [41,76], involves alterations in gene expression in the mesocorticolimbic systems. Dopamine-dependent rewarding attenuates the overall excitability of GABAergic neurons, and these processes are exhibited in decreased expression of all *Slc\** genes encoding transporters of amino acids, glucose, nucleoside sugars, neurotransmitters, vesicular transporters, inorganic and organic ions, which leads to changes in signaling cascades through reduced activation of GABAergic MSNs in the NAcc, and thereby to decreased inhibitory control of aggression [74].

However, according to available data, no contemporary drugs have been developed specifically to activate SLC transporter activity. It is very important to find new targets for correction of metabolic and neurotransmitter processes implicated in pathological states. Altered expression of the genes with the largest number of correlations with other gene expression may serve as a prognostic target and a tool to search for new generation drugs.

We have previously found downregulation of collagen genes encoding proteins that are the main components of the extracellular matrix in the ventral segmental areas of male mice with repeated experiences of aggression [77]. These data, together with decreased expression of *Slc \** genes, support the view that neural plasticity accompanied by a decreased synaptic connections in brain regions that are involved in neuropsychiatric disorders, may be reduced. This leads to a decrease in cell survival, as well as to a decrease in the efficiency of neural connections. We hope our research will help find a new approach to restore neural plasticity which plays an important role in development of mood disorders.

## Abbreviations

MSN: Medium Spiny Neurons;
*Slc\**: solute carrier family;
PCA: Principal Component Analysis;
AHC: Agglomerative Hierarchical Clustering;
FPKM: fragments per kilobase of transcript per million mapped reads;
VTA: ventral tegmental area;
NAcc: nucleus accumbens;
PFC: prefrontal cortex.
W: winners;
C: controls.

## Ethics approval

All procedures were in compliance with the European Communities Council Directive 210/63/EU on September 22, 2010. The study was approved by the Bioethical Commission [Scientific Council No9) in the Institute of Cytology and Genetics SD RAS of March, 24, 2010, N 613 [Novosibirsk).

## Consent for publication

N/A

## Acknowledgement

We thank O.A. Kharlamova for stylistic assistance in writing the manuscript. Preparation and maintenance of experimental animals was carried out in the vivarium of the Institute of Cytology and Genetics SB RAS and was funded under ICG SB RAS, BP No 0324-2019-0041-C-01. This work was supported by Russian Science Foundation (grant No19-15-00026) (to NNK). There is none role of the funding body in the design of the study and collection, analysis, and interpretation of data and in writing the manuscript.

## Availability of data and materials

The additional statistics of data obtained used to support the findings of this study are available from Supplement 1 (Tables S1, differentially expressed *Slc\** genes in FPKM units) and are cited at relevant places within the text. The other datasets generated during the current study are available from the corresponding author on reasonable request.

## Authors’ information

DAS, VNB, ILK, AGG, OER, NNK: Laboratory of Neuropathology Modeling; NNK, DAS, ILK, AGG: Neurogenetics of Social Behavior Sector; The FRC Institute of Cytology and Genetics SB RAS, Novosibirsk, Russia DAS, ILK, AGG contributed substantially to behavioral data acquisition, received brain materials; DAS, VNB and OER implemented bioinformatics assessing of RNA-Seq data, analyzed and interpreted data. NNK performed study design, analyzed the RNA-seq database and interpreted data, wrote the main manuscript text. All authors gave final approval.

The authors declare that they have no competing interests.

**Supplement 1, Table S1**. Samples expression level (FPKM) of differentially expressed *Slc\** genes in the VTA, NAcc and PFC.

**Supplement 1, Table S2**. AHC of the *Slc*\* genes encoding transcripts in the VTA, NAcc and PFC.

**Supplement 1, Table S3**. Correlation coefficients between differentially expressed *Slc*\* genes in the VTA, NAcc and PFC.

## References

1. https://en.wikipedia.org/wiki/Solute_carrier_family

2. http://slc.bioparadigms.org/

3. https://en.wikipedia.org/wiki/Transporter_Classification_Database

4. He L, Vasiliou K, Nebert DW (2009): Analysis and update of the human solute carrier (SLC) gene superfamily. Hum Genomics 3(2):195–206.

5. César-Razquin A, Snijder B, Frappier-Brinton T, Isserlin R, Gyimesi G, Bai X, et al. (2015): Call for Systematic Research on Solute Carriers. Cell 162: 478–487.

6. Saier MH Jr, Reddy VS, Tsu BV, Ahmed MS, Li C, Moreno-Hagelsieb G (2016): The transporter classification database (TCDB): recent advances. Nucleic Acids Res 44(D): D372–D379.

7. Lin L, Yee SW, Kim RB, Giacomini KM (2015): SLC Transporters as therapeutic targets: emerging opportunities. Nat Rev Drug Discov 14(8): 543–560.

8. Jankovic J, Clarence-Smith K (2011): Tetrabenazine for the treatment of chorea and other hyperkinetic movement disorders. Expert Rev Neurother 11: 1509–1523.

9. Nickell JR, Siripurapu KB, Vartak A, Crooks PA, Dwoskin LP (2014): The vesicular monoamine transporter-2: an important pharmacological target for the discovery of novel therapeutics to treat methamphetamine abuse. Adv Pharmacol 69: 71–106.

10. Harvey RJ, Yee BK (2013): Glycine transporters as novel therapeutic targets in schizophrenia, alcohol dependence and pain. Nat Rev Drug Discov 12: 866–885.

11. Kudryavtseva NN, Bondar NP, Avgustinovich DF (2002): Association between experience of aggression and anxiety in male mice. Behav Brain Res 133(1): 83–93.

12. Kudryavtseva NN (2006): Psychopathology of repeated aggression: a neurobiological aspect. In: Morgan JP, editors. Perspectives on the Psychology of Aggression. NOVA Science Publishers, pp 35–64.

13. Kudryavtseva NN, Smagin DA, Kovalenko IL, Vishnivetskaya GB (2014): Repeated positive fighting experience in male inbred mice. Nat Prot 9:11: 2705-2717.

14. Kovalenko IL, Smagin DA, Galyamina AG, Kudryavtseva NN (2015): Hyperactivity and abnormal exploratory activity developing in the CD-1 male mice under chronic experience of aggression and social defeats in daily agonistic interactions. J Behav Brain Sci 5(11): 478–490.

15. Smagin DA, Galyamina AG, Kovalenko I L, Babenko VN, Tamkovich NV, Borisov SA, et al. (2018): Differentially expressed neurotransmitter genes in the dorsal striatum of male mice with psychomotor disturbances. Zh Vyssh Nerv Deiat Im I P Pavlova 68(2): 227–249.

16. Kudryavtseva NN (2020): Positive fighting experience, addiction-like state, and relapse: Retrospective analysis of experimental studies. ZAggress Violent Behav 52, (5-6), 101403.

17. Filipenko ML, Alekseyenko OV, Beilina AG, Kamynina TP, Kudryavtseva NN (2001): Increase of tyrosine hydroxylase and dopamine transporter mRNA levels in ventral tegmental area of male mice under influence of repeated aggression experience. Brain Res, Mol Brain Res 96: 77–81.

18. Filipenko ML, Beylina AG, Alekseyenko OV, Timofeeva OA, Avgustinovich DF, Kudryavtseva NN (2002): Association between brain COMT gene expression and aggressive experience in daily agonistic confrontations in male mice. In: McCarty R, Aguilera G, Sabban E, Kvetnyansky R, editors. Stress: Neural, Endocrine and Molecular Studies. New York: Taylor & Francis, pp 157–161.

19. Bondar NP, Boyarskikh UA, Kovalenko IL, Filipenko ML, Kudryavtseva NN (2009): Molecular implications of repeated aggression: Th, Dat1, Snca and Bdnf gene expression in the ventral teg mental area of victorious male mice. PLoS One 4(1): e4190.

20. Kudryavtseva NN, Filipenko ML, Bakshtanovskaya IV, Avgustinovich DF, Alekseenko OV, Beilina AG (2004): Changes in the expression of monoaminergic genes under the influence of repeated experience of agonistic interactions: From behavior to gene. Russ J Genet 40(6): 590–604.

21. Smagin DA, Boyarskikh UA, Bondar NP, Filipenko ML, Kudryavtseva NN (2013): Reduction of serotonergic gene expression in the midbrain raphe nuclei under positive fighting experience. Adv Biosci Biotech 4(10B): 36–44.

22. Kudriavtseva NN, Bakshtanovskaia IV (1991): The neurochemical control of aggression and submission. Zh Vyssh Nerv Deiat Im I P Pavlova 41(3): 459–466.

23. Schultz W (1998): Predictive reward signal of dopamine neurons. J Neurophysiol 80(1): 1–27.

24. Dayan P, Balleine BW (2002): Reward, motivation, and reinforcement learning. Neuron 36(2): 285–298.

25. Wise RA (2004): Dopamine, learning and motivation. Nat Rev Neurosci 5(6): 483–494

26. Berridge KC (2007): The debate over dopamine’s role in reward: the case for incentive salience. Psychopharmacology 191(3): 391–431.

27. Ikemoto S (2007): Dopamine reward circuitry: two projection systems from the ventral midbrain to the nucleus accumbens-olfactory tubercle complex. Brain Res Rev 56(1): 27–78.

28. Russo SJ, Nestler EJ (2013): The brain reward circuitry in mood disorders. Nat Rev Neurosci 14 (9): 609–625.

29. Aleyasin H, Flanigan ME, Russo SJ (2018): Neurocircuitry of aggression and aggression seeking behavior: nose poking into brain circuitry controlling aggression. Curr Opin Neurobiol 49: 184–191.

30. Kudryavtseva NN (1991): The sensory contact model for the study of aggressive and submissive behaviors in male mice. Aggress Behav 17(5): 285–291.

31. http://mouse.brain-map.org/static/atlas

32. Margolis EB, Lock H, Hjelmstad GO, Fields HL (2006): The ventral tegmental area revisited: is there an electrophysiological marker for dopaminergic neurons? J Physiol 577 (Pt 3): 907–924.

33. Sesack SR, Grace AA (2010): Cortico-Basal Ganglia reward network: microcircuitry. Neuropsychopharmacology 35(1): 27–47.

34. Trutti AC, Mulder MJ, Hommel B, Forstmann BU (2019): Functional neuroanatomical review of the ventral tegmental. Neuroimage 191:258–268.

35. Walsh JJ, Han MH (2014): The heterogeneity of ventral tegmental area neurons: Projection functions in a mood related context. Neuroscience 282: 101–108.

36. Morales M, Margolis EB (2017): Ventral tegmental area: cellular heterogeneity, connectivity and behaviour. Nat Rev Neurosci 18(2), 73–85.

37. Bourdy R, Barrot M (2012): A new control center for dopaminergic systems: pulling the VTA by the tail. Trends Neurosci 35 (11): 681–690.

38. Nair–Roberts RG, Chatelain–Badie SD, Benson E, White–Cooper H, Bolam JP, Ungless MA (2008): Stereological estimates of dopaminergic, GABAergic and glutamatergic neurons in the ventral tegmental area, substantia nigra and retrorubral field in the rat. Neuroscience 152: 1024–1031.

39. Yamaguchi T, Sheen W, Morales M (2007): Glutamatergic neurons are present in the rat ventral tegmental area. Eur J Neurosci 25: 106–118.

40. https://en.wikipedia.org/wiki/Nucleus_accumbens#cite_note-45

41. Robinson AJ, Nestler EJ [2011): Transcriptional and epigenetic mechanisms of addiction. Nat Rev Neurosci 12(11): 623–637.

42. Meredith GE, Pennartz CM, Groenewegen HJ (1993): The cellular framework for chemical signalling in the nucleus accumbens. Prog Brain Res 99: 3–24.

43. Olsen CM (2011): Natural rewards, neuroplasticity, and non-drug addictions. Neuropharmacology. 61(7): 1109–1122.

44. Alvarez JA, Emory E (2006): Executive function and the frontal lobes: a meta-analytic review. Neuropsychol Rev 16 (1): 17–42.

45. Steketee JD (2003): Neurotransmitter systems of the medial prefrontal cortex: potential role in sensitization to psychostimulants. Brain Res Rev 41(2-3): 203–228.

46. Del Arco A, Mora F (2009): Neurotransmitters and prefrontal cortex-limbic system interactions: implications for plasticity and psychiatric disorders. J Neural Transm [Vienna) 116(8): 941–952.

47. Trapnell C, Hendrickson DG, Sauvageau M, Goff L, Rinn JL, Pachter L (2013): Differential analysis of gene regulation at transcript resolution with RNA-seq. ZNat Biotechnol 31: 46–53.

48. Dahlin A, Royall J, Hohmann JG, Wang J (2009): Expression profiling of the solute carrier gene family in the mouse brain. J Pharmacol Exp Ther 329(2): 558–570.

49. Babenko VN, Smagin DA, Galyamina AG, Kovalenko IL, Kudryavtseva NN (2018): Altered Slc25 family gene expression as markers of mitochondrial dysfunction in brain regions under experimental mixed anxiety/depression-like disorder. BMC Neurosci 19(1): 79.

50. Babenko VN, Galyamina AG, Rogozin IB, Smagin DA, Kudryavtseva NN (2020a): Dopamine response gene pathways in dorsal striatum MSNs from a gene expression viewpoint: cAMP-mediated gene networks. BMC Neurosci 21(1): 12.

51. Babenko VN, Smagin DA, Galyamina AG, Kovalenko IL, Kudryavtseva NN (2020b): Differentially expressed genes of the Slc6a family as markers of altered brain neurotransmitter system function in pathological state in mice. Neurosci Behav Physiol 50: 199–209.

52. Fields HL, Hjelmstad GO, Margolis EB, Nicola SM (2007): Ventral tegmental area neurons in learned appetitive behavior and positive reinforcement. Annu Rev Neurosci 30: 289–316.

53. Grace AA (1991): Phasic versus tonic dopamine release and the modulation of dopamine system responsivity: a hypothesis for the etiology of schizophrenia. Neuroscience 41(1): 1–24.

54. Tomasi D, Volkow ND (2014): Functional connectivity of substantia nigra and ventral tegmental area: maturation during adolescence and effects of ADHD. Cereb Cortex 24 (4): 935–944.

55. Barker DJ, Root DH, Zhang S, Morales M (2016): Multiplexed neurochemical signaling by neurons of the ventral tegmental area. ZJ Chem Neuroanat 73: 33–42.

56. Polter AM, Kauer JA (2014): Stress and VTA synapses: implications for addiction and depression. Eur J Neurosci 39(7): 1179–1188.

57. Adinoff B (2004): Neurobiologic processes in drug reward and addiction. Harvard Rev Psychiat 12(6): 305–320.

58. Juarez B, Han MH (2016): Diversity of dopaminergic neural circuits in response to drug exposure. Neuropsychopharmacology 41,10: 2424–2446

59. Niehaus JL, Cruz-Bermudez ND, Kauer JA (2009): Plasticity of addiction: a mesolimbic dopamine short-circuit? Am J Addict 18(4): 259–271.

60. https://en.wikipedia.org/wiki/Tyrosine_hydroxylase

61. Miczek KA, Faccidomo SP, Fish EW, DeBold JF (2007): Neurochemistry and molecular neurobiology of aggressive behavior. In: Lajtha A, Blaustein JD, editors. Handbook of Neurochemistry and Molecular Neurobiology: Behavioral Neurochemistry, Verlag Berlin Heidelberg: Springer, pp 285–336.

62. Van Erp AM, Miczek KA (2000): Aggressive behavior, increased accumbal dopamine, and decreased cortical serotonin in rats. J Neurosci 20: 9320–9325.

63. Kudryavtseva NN (2007): Straub tail, the deprivation effect and addiction to aggression. In: O’Neal PW, editors. Motivation of Health Behavior. NOVA Science Publishers, pp 97–110

64. Kudryavtseva NN, Smagin DA, Bondar NP (2011): Modeling fighting deprivation effect in mouse repeated aggression paradigm. Progr Neuropsychopharmacol Biol Psychiatry 35(6): 1472–1478.

65. Kudryavtseva NN, Lipina TV, Koryakina LA (1999): Effects of haloperidol on communicative and aggressive behavior in male mice with different experiences of aggression. Pharmacol Biochem Behav 63(2): 229–236.

66. Moyer KE (1987): Violence and aggression. New-York: Paragon House.

67. Scott JP (1966): Agonistic behavior of mice and rats: a review. Am Zool 6: 4: 683-701.

68. Scott JP (1971): Theoretical issues concerning the origin and causes of fighting. In Eleftheriou BE, Scott JP, editors. The Physiology of Aggression and Defeat. New-York: Plenum, pp 11–42.

69. Brain PF (1979): The adaptiveness of house mouse aggression. In Brain PF, Mainardi D, Parmigiani S, editors. House Mouse Aggression. A Model for Understanding the Evolution of Social Behavior. Chur: Harwood Academic Publishers, pp 1–21.

70. Ramirez JM, Bonniot-Cabanac MC, Cabanac M (2005): Can aggression provide pleasure? Eur Psychol 10: 2: 136-145.

71. Kudryavtseva NN (2000): An experimental approach to the study of learned aggression. Aggress Behav 26(3): 241–256.

72. Golden SA, Shaham Y (2018): Aggression, addiction and relapse: A new frontier in psychiatry. Neuropsychopharmacology 43(1): 224–225.

73. Golden SA, Jin M, Shaham Y (2019): Animal models of (or for) aggression reward, addiction, and relapse: behavior and circuits. J Neurosci 39 (21): 3996–4008.

74. Carlezon WA, Thomas MJ (2009): Biological substrates of reward and aversion: a nucleus accumbens activity hypothesis. Neuropharmacology 56 (Suppl 1): 122–132.

75. Day JJ, Carelli RM (2007): The nucleus accumbens and Pavlovian reward learning. Neuroscientist. 13 (2): 148–159.

76. Steiner H, Van Waes V (2013): Addiction-related gene regulation: risks of exposure to cognitive enhancers vs. other psychostimulants. Prog Neurobiol 100: 60–80.

77. Smagin DA, Galyamina AG, Kovalenko IL, Babenko VN, Kudryavtseva NN (2019) Aberrant expression of collagen gene family in the brain regions of male mice with behavioral psychopathologies induced by chronic agonistic interactions. Biomed Res Int 2019:7276389, 13 pages. doi: 10.1155/2019/7276389.

